# *SimPhy*: Phylogenomic Simulation of Gene, Locus and Species Trees

**DOI:** 10.1101/021709

**Authors:** Diego Mallo, Leonardo de Oliveira Martins, David Posada

## Abstract

We present here a fast and flexible software –*SimPhy*– for the simulation of multiple gene families evolving under incomplete lineage sorting, gene duplication and loss, horizontal gene transfer –all three potentially leading to the species tree/gene tree discordance– and gene conversion. *SimPhy* implements a hierarchical phylogenetic model in which the evolution of species, locus and gene trees is governed by global and local parameters (e.g., genome-wide, species-specific, locus-specific), that can be fixed or be sampled from *a priori* statistical distributions. *SimPhy* also incorporates comprehensive models of substitution rate variation among lineages (uncorrelated relaxed clocks) and the capability of simulating partitioned nucleotide, codon and protein multilocus sequence alignments under a plethora of substitution models using the program INDELible. We validate *SimPhy*’s output using theoretical expectations and other programs, and show that it scales extremely well with complex models and/or large trees, being an order of magnitude faster than the most similar program (DLCoal-Sim). In addition, we demonstrate how *SimPhy* can be useful to understand interactions among different evolutionary processes, conducting a simulation study to characterize the systematic overestimation of the duplication time when using standard reconciliation methods. *SimPhy* is available at https://github.com/adamallo/SimPhy, where users can find the source code, pre-compiled executables, a detailed manual and example cases.

---

Recent advances in sequencing technologies have furnished the expansion of genome-wide phylogenetic studies, unveiling not only extensive phylogenomic incongruence (Jeffroy, Brinkmann, et al. 2006; Salichos and Rokas 2013) but also bringing back to the spotlight the consideration of how ancestral polymorphisms sort within populations (Edwards 2009). Indeed, it is well known that gene and species phylogenies can be inconsistent due to evolutionary processes like incomplete lineage sorting (ILS), gene duplication and loss (GDL) and horizontal gene transfer (HGT) (Goodman, Czelusniak, et al. 1979; Pamilo and Nei 1988; Takahata 1989; Maddison 1997; Page and Charleston 1997). Not surprisingly, the gene tree / species tree ‘dilemma’ has received a lot of attention in recent years, and consequently a plethora of species tree reconstruction methods have been published (Chaudhary, Bansal, et al. 2010; e.g., Heled and Drummond 2010; Liu, Yu, et al. 2010; Boussau, Szollosi, et al. 2013; De Oliveira Martins, Mallo, et al. 2014; Mirarab and Warnow 2015; see Mallo, de Oliveira Martins, et al. 2014a for a review). Although several benchmarks of species tree methods have been carried out (Leache and Rannala 2011; Yang and Warnow 2011; Bayzid and Warnow 2013; Mirarab, Bayzid, et al. 2014) they have generally focused on single causes of phylogenetic discordance. Nevertheless, ILS, GDL and HGT can act simultaneously during genome evolution yielding synergistic evolutionary scenarios (Mallo, De Oliveira Martins, et al. 2014b) and therefore it would be convenient to consider them all together. However, none of the state-of-the-art simulation programs is able to yield scenarios that consider these processes at once. Rather, they usually focus on single evolutionary processes, which partially explains why benchmarking studies have usually been restrictive in this regard. Thus, ILS is usually simulated with tools that implement different extensions of the multispecies coalescent model (MSC) (Rannala and Yang 2003), like Mesquite (Maddison and Maddison 2015), MCcoal (Rannala and Yang 2003) or GUMS (Heled, Bryant, et al. 2013). Moreover, for small species trees, structured coalescent simulators like ms (Hudson 2002), SGWE (Arenas and Posada 2014) or scrm (Staab, Zhu, et al. 2015) could be also used. Apart from standalone programs, there are also phylogenetic libraries like DendroPy (Sukumaran and Holder 2010) that include the simulation of ILS. GDL can be modeled using birth-death processes traversing the species tree like in Arvestad, Berglund, et al. (2003), and HGT is usually simulated as a Poisson distributed series of transfer events, like in HGT_simul (Galtier 2007). Nowadays, only very few tools are able to simulate phylogenies jointly considering multiple sources of phylogenomic incongruence, like PrIME-GenPhyloData (Sjostrand, Arvestad, et al. 2013) and DLCoal_sim (Rasmussen and Kellis 2012). The former combines GDL and HGT, while the later considers GDL and ILS. In addition, there are also genome simulators like ALF-A (Dalquen, Anisimova, et al. 2012) or EvolSimulator (Beiko and Charlebois 2007), that can integrate several evolutionary processes at the genomic level but do not consider population level events, like ILS. Altogether, we are not aware of any tool able to simulate phylogenetic trees considering the joint action of ILS, GDL and HGT.

In order to facilitate more realistic simulations, we present here a fast and flexible simulation tool –*SimPhy*– that can simulate the evolution of multiple gene families under ILS, GDL, HGT and gene conversion (GC). *SimPhy* implements a flexible hierarchical parameterization scheme that considers genome-wide and gene family specific conditions, including different sources for evolutionary rate variation among lineages. Moreover, these parameters can be fixed or sampled from statistical distributions defined by the user. In addition, *SimPhy* is able to produce multilocus sequence alignment using INDELible (Fletcher and Yang 2009). All these features make *SimPhy* a powerful tool to understand the interaction among ILS, GDL and HGT or for a comprehensive benchmarking of phylogenomic methods.

## SIMULATION OF GENE, LOCUS AND SPECIES TREES WITH SIMPHY

*SimPhy* simulates the evolution of multiple gene families under a hierarchical phylogenomic model in which gene trees evolve inside locus trees, which in turn evolve along a single species tree (Fig. 1). While this three-tree model has already been implemented in the program DLCoal_sim (Rasmussen and Kellis 2012), *SimPhy* extends this approach in multiple ways. Apart from ILS and GDL, *SimPhy* also jointly considers HGT and GC, plus species extinction. Furthermore, GDL, HGT and GC rates are allowed to vary among gene families. *SimPhy* also relaxes the assumption of a strict molecular clock and implements different sources of rate heterogeneity among lineages at the species, gene family and gene tree level. Moreover, the generation time is allowed to vary along the species tree, incorporating an additional layer of rate variation. *SimPhy*’s simulation parameters can be sampled from user-specified distributions at the species, locus and gene tree level, and in some cases made interdependent. This allows users to carry out simulations following a very flexible strategy in which each parameter is defined by a prior distribution that can represent specific, biologically relevant scenarios. Nevertheless, (classic) simulation studies based on combinations of fixed, discrete parameter values can also be easily implemented.

**Figure 1.**
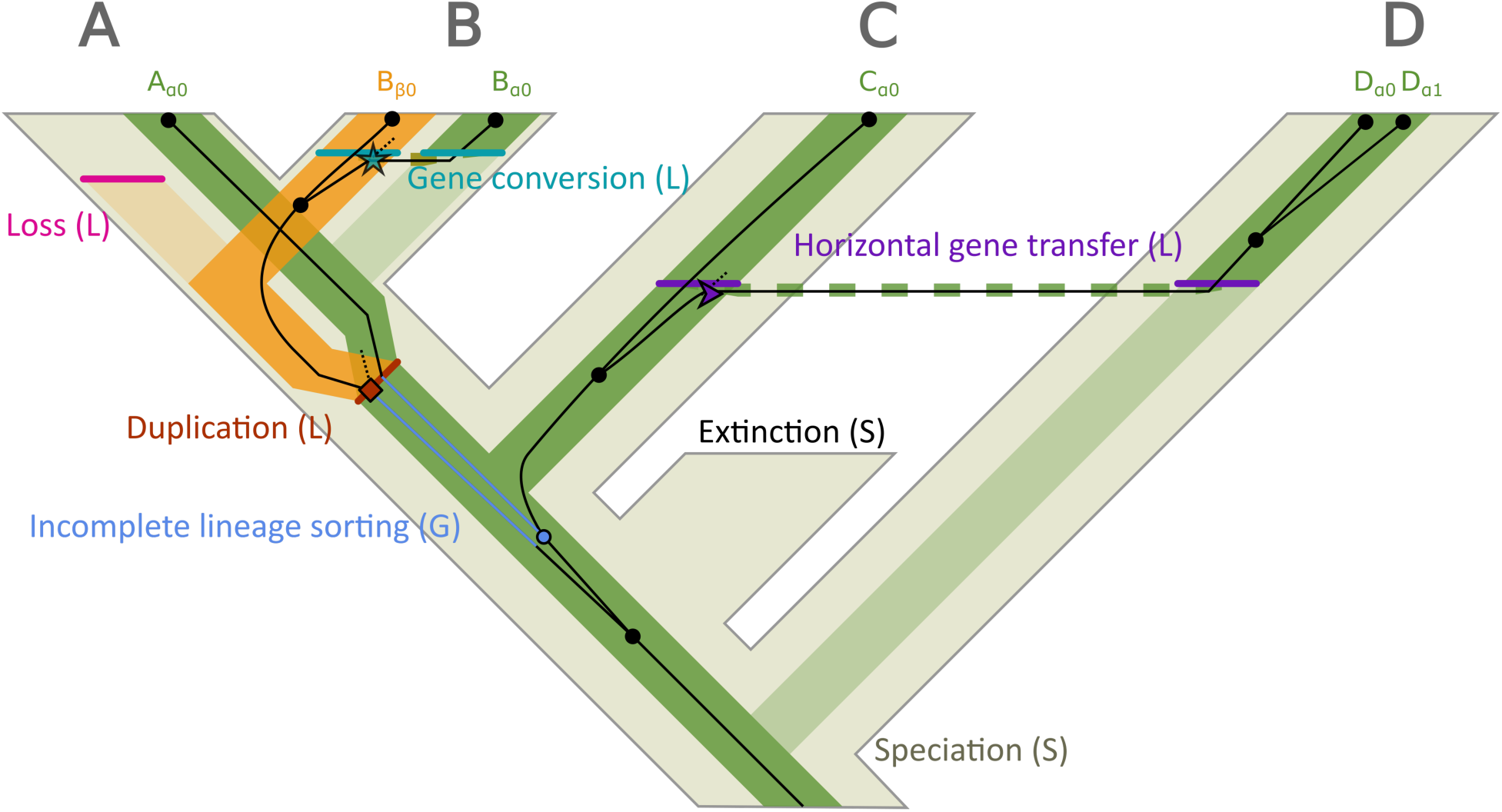
Three-tree model and evolutionary events simulated in *SimPhy*. The figure shows a species tree (thick light tree in the background) that embeds a locus tree (medium-thick lines) that includes a gene tree (thin dark lines with nodes represented by black dots). The sample consists of six gene copies (A_α0_, B_β0_, B_α0_, C_α0_, D_α0_ and D_α1_) that belong to two loci (α and β) in four different species (A, B, C and D) and five individuals (A_0_, B_0_, C_0_, D_0_ and D_1_). Letters in parenthesis indicate at which phylogenetic level an event takes place (S= species tree, L= locus tree, G= gene tree). Coalescent bounds generated by GDL, HGT or GC events are indicated with an icon (duplication: square, horizontal gene transfer: arrow, gene conversion: star). Dashed lines in the locus tree, generated by HGT and GC, link locus tree branches that are depicted apart for clarity. Lighter locus tree branches represent lost branches due to gene loss or locus replacement by HGT or GC. Dashed gene tree branches indicate lost/unsampled lineages in the original locus from the individual where the duplication took place.

### The Three-tree Model

*SimPhy* implements and extends a hierarchical phylogenetic model (Rasmussen and Kellis 2012) that considers three different layers: species, locus and gene trees.

#### Species trees

Depict the evolutionary history of the sampled organisms. Species tree nodes represent speciation events, while the branches reflect population history: branch length and width represent elapsed time and effective population size (Ne), respectively. Note that we consider species as a diverging interbreeding structure regardless of any taxonomic rank. Therefore, species trees might be equivalent to population trees when the organismal units of interest are conspecific populations.

#### Locus trees

Represent the evolutionary history of the sampled loci for a given gene family. Since the loci exist inside individuals evolving as part of a population, the locus tree is embedded within the species tree. In a locus tree the nodes depict either genetic divergence due to speciation in the embedding species tree or locus-level events like duplication, loss, horizontal gene transfer or gene conversions, while branch lengths and widths represent time and Ne as before.

#### Gene trees

Represent the evolutionary history of the sampled gene copies, which evolve inside a locus tree. Gene tree nodes indicate coalescent events, which looking forward in time correspond to the process of DNA replication and divergence, and which in the absence of migration can only occur before the speciation time. The lengths of the gene tree branches represent the expected number of substitutions per site.

### Evolutionary Processes Implemented

*SimPhy*’s generative model encompasses different evolutionary processes:

1. **Speciation**.– Separation of one ancestral species into two new species that do not interbreed. Speciation events are represented by nodes in the species tree, and can also be reflected in the locus tree.
2. **Extinction**.– Disappearance of a species. Extinctions take place during the simulation of the species tree, affecting its final topology, but are not considered further.
3. **Gene duplication**.– Copy of one gene into a new locus in every individual of the population (i.e., we assume no duplication polymorphism). Gene duplication events generate nodes in the locus tree.
4. **Gene loss**.– Deletion of one gene in every individual of the population (i.e., we assume no loss polymorphism). Losses generate locus tree leaves that do not reach the present, and that are not considered during the gene tree simulation process.
5. **Horizontal gene transfer**.– Copy of one locus from one species to another contemporary species via replacement. The transfer affects every individual in the receptor species (i.e., there is no transfer polymorphism). Each transfer generates two nodes in the locus tree, one representing the loss of the replaced locus (receptor) and another showing the incorporation of the transferred lineage.
6. **Gene conversion**.– Replacement of one homolog by another within a single species. This conversion affects every individual in that species (i.e., there is no gene conversion polymorphism). Each gene conversion generates two nodes in the locus tree, one representing the loss of the replaced locus (receptor) and another showing the incorporation of the converted lineage.
7. **Lineage sorting**.– Consideration of the coalescent history of the sampled gene copies, allowing their history to be incompatible with the species tree history. It is implicitly reflected at the gene tree level, and can be spotted when mapping locus and gene tree nodes.

These evolutionary processes are modeled by *SimPhy* using a mixture of existing and original strategies. The **species tree** is simulated (or user-defined, see *Simulation process*) by either a Yule (Yule 1925) or a birth-death (Kendall 1948) process, considering speciations and extinctions. At this point, *SimPhy* only considers extant species. Each **locus tree** is described by a birth-death process that considers duplication and losses, coupled with other two pure birth processes that describe the HGT and GC events. Regarding the last two, *SimPhy* incorporates two variants, one in which the receptor is randomly chosen from the candidates and another that takes into account the evolutionary distance between donors and candidate receptors – i.e., with reception probability inversely proportional to the phylogenetic distance.

Duplications, horizontal gene transfers and gene conversions act as coalescent bounds for the subtree that corresponds to the new/replaced locus, since these events initially affect a single individual and only afterwards become fixed in the population (Fig. 2). Nevertheless, *SimPhy* does not consider locus polymorphism (i.e., every individual in a particular lineage carries the same number of gene copies for a given gene family), assuming that these events either get fixed or drifted away in all the descendant lineages. **Gene trees** are modeled by the multilocus coalescent model (Rasmussen and Kellis 2012) expanded to consider bounded subtrees generated by HGT and GC. This model is built upon the multispecies coalescent model (Rannala and Yang 2003) and therefore inherits the same assumptions: each branch of the container tree is composed by a perfect Wright-Fisher population (constant effective population size, non-overlapping generations, random mating, neutrality), no recombination within loci and free recombination between loci. Consequently, all the evolutionary processes except speciation and extinction are considered independently for each gene family. Some of these assumptions might be relaxed in future versions of *SimPhy*, for example allowing migration of individuals between species and or recombination within loci.

**Figure 2.**
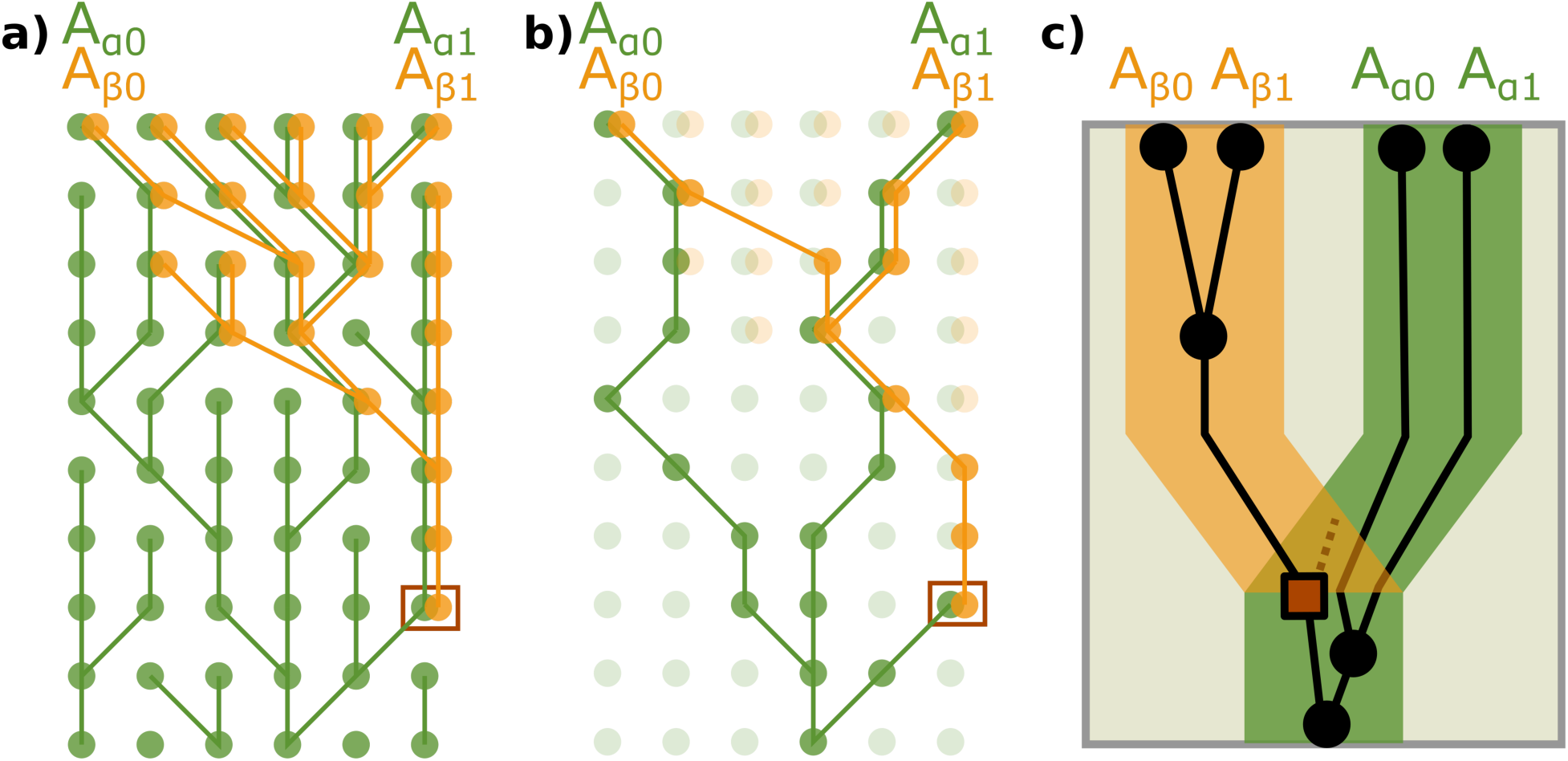
Coalescent bound enforced by duplication. The figure represents the evolutionary relationships of four gene copies (A_β0_, A_β1_, A_α0_ and A_α1_) pertaining to two individuals (A0 and A1) of the same species (A) with two loci (α and β). The locus β was originated by duplication in the individual surrounded by a square. We assume that the duplication occurs initially in one individual, and afterwards spreads throughout the whole population. Thus, lineages coming from the β lineage cannot exist below the duplication event, imposing a coalescent bound (i.e., they have to coalesce before the duplication event -above in this figure). The three subfigures represent alternative views of the same scenario. Subfigure a) depicts the relationships of the whole population, while in subfigure b) only the sampled lineages are highlighted. Finally, subfigure c) shows the three-tree representation of the sample genealogy.

### Simulation Process

#### Species trees

Sampled either using a pure birth (Yule) model or a birth-death model, parameterized by birth (speciation) and death (extinction) rates (given in number of events per generation) and either number of leaves, tree height or both. The simple sampling approach algorithm (SSA) is used when the number of leaves is not specified, while the birth-death rate sampling approach (BDSA) algorithm is used otherwise (see Hartmann, Wong, et al. 2010 for more details). An outgroup species can be added to the resulting species tree before the simulation of the locus trees.

#### Locus trees

Simulated using a SSA to sample a birth-death process –describing duplications and losses– coupled with two additional pure birth processes –describing horizontal gene transfers and gene conversions– along each species tree branch, in a preorder fashion. This is followed by a second preorder traversal that samples receptors from contemporary candidates for each HGT/GC, completing the appropriate SPR branch rearrangement in order to generate the definitive locus tree.

#### Gene trees

Sampled traversing the locus tree in post-order. Locus tree branches that not pertain to bounded subtrees (i.e., representing the existing locus before any bounding duplication/HGT/GC event) are modeled by the multispecies coalescent, and therefore sampled using a rejection sampling strategy where waiting times come from an Exponential distribution. However, simulation along branches pertaining to bounded locus subtrees implies a much more complex strategy, based on sampling the number of gene tree lineages going through every node of the locus tree, and obtaining the coalescent times conditioned on these numbers. This approach requires the calculation of lineage count probabilities for every branch of the these subtrees, for which *SimPhy* uses a dynamic programing algorithm modified from the one proposed by Rasmussen and Kellis (2012). A preorder traversal is afterwards conducted to sample the number of lineage counts going across every locus subtree branch, using the inverse transform sampling method with the CDF of the number of input lineages conditioned on the pre-calculated input probabilities, output lineage counts, effective population size and branch length for every branch of the considered subtree (Appendix 1). This procedure starts at the root of each bounded subtree, where the number of output lineages is fixed to one –because of the bound–, and then samples the counts along the tree, stopping at the leaves. With the lineage counts already set, the coalescent times are sampled using the inverse transform of the CDF of the coalescent times conditioned on lineage counts (as in Rasmussen and Kellis 2012).

### Distribution-driven Parameterization

While most simulation studies are parameterized using a grid consisting of combinations of discrete parameter values, *SimPhy* has the capability of sampling parameter values from prior statistical distributions (as in Darriba, Taboada, et al. 2012; see also Leigh and Bryant 2015). Such distributions are defined by the user, and currently include Uniform, Normal, Lognormal, Exponential and Gamma plus the possibility of fixing parameter values. Different parameters are sampled through the different simulation layers (i.e., for each species, locus or gene tree), with the possibility of specifying certain dependencies among parameters (i.e., with some parameters acting as hyper or hyper-hyper-parameters) (Fig. 3).

**Figure 3.**
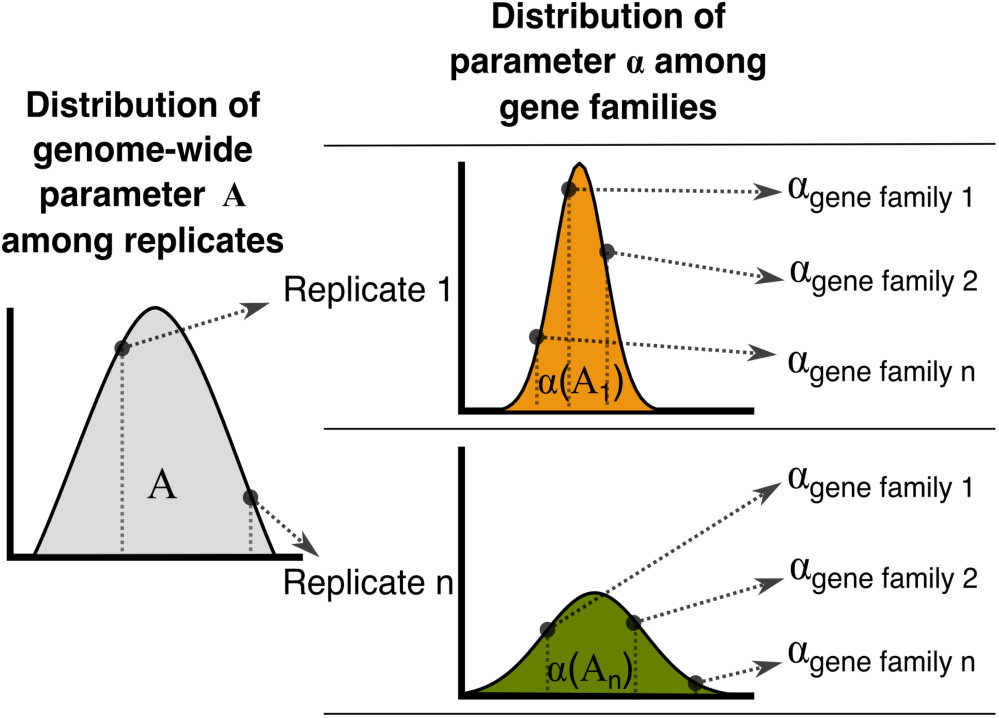
Distribution-driven parameterization. For every simulation replicate parameter A is sampled from a given distribution. The sampled value of A will shape the distribution among gene families of our parameter of interest, α, for that replicate (i.e., A acts a hyperparameter for α). Then, for every gene family the α parameter is sampled from the resulting distribution in order to determine its evolution. This particular schema can be applied to all gene-family-specific parameters like duplication, loss, horizontal gene transfer and gene conversion rates. Similar schemas with more or less layers can be applied to other simulation parameters. Different statistical distributions can be specified by the user (e.g., Uniform, Normal, Exponential, Gamma).

Under the standard simulation workflow, *SimPhy* samples for each replicate genome-wide parameters, species tree parameters, species-specific and gene-family-specific rate variation parameters, and number of gene families. Genome-wide parameters control the expected distribution of duplications, losses, horizontal gene transfers and gene conversion events across gene families. Species tree parameters include speciation and extinction rates, species tree height, number of taxa, relative distance of the outgroup, number of individuals per species, effective population size, substitution rate and generation time. For each gene family, specific duplication, loss, horizontal gene transfer and gene conversion rates are sampled from the distributions specified at the genome-wide level. The variation of the substitution rate across is also specified using statistical distributions at different levels, as detailed in the next section.

### Substitution Rate Variation

The generative model of *SimPhy*, being based on coalescent theory and birth-death models, is in principle intrinsically ultrametric and therefore follows a strict molecular clock. However, it is well known that many real data sets deviate from a strict molecular clock (see Ho, Lanfear, et al. 2011). We have therefore implemented in *SimPhy* different sources of rate variation among lineages –lineage specific, gene family specific and gene-by-lineage specific (Fig. 4)– which modify the branches of the gene trees. In all cases a rate multiplier is sampled from a Gamma distribution, whose mean is forced to be one in order to preserve the mean substitution rate, and that therefore can be parameterized by a single *alpha* shape parameter. Since these multipliers are independently sampled, this approach is equivalent to the use of uncorrelated relaxed clocks (Drummond, Ho, et al. 2006).

**Figure 4.**
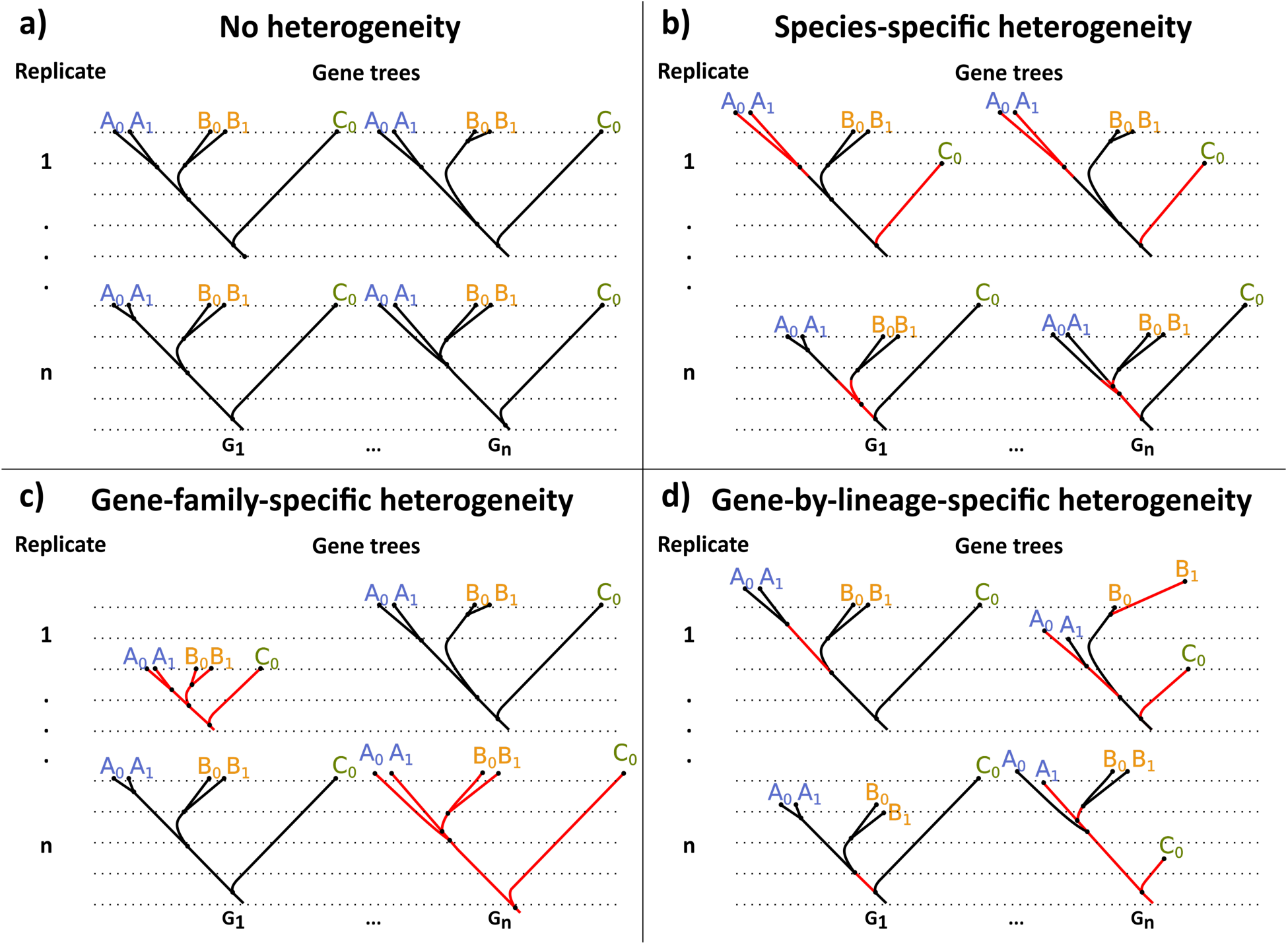
Sources of substitution rate heterogeneity among lineages available in *SimPhy*. Each subfigure represents the same simulation scenario but with different types of rate variation. Each simulation scenario consists of two independent replicates (rows) of two gene families (columns) in three species (A,B,C) with five individuals (A0,A1,B0,B1,C0). Subfigure a) shows trees simulated without substitution rate heterogeneity –hence ultrametric– with topologies and branch lengths generated by the coalescent process. Subfigure b) depicts a case of species-specific heterogeneity, where changes in rate affect whole ancestral/contemporary species genome-wide (e.g., for every gene family in replicate 1 species A branches evolve faster and species C branches slower). Subfigure c) is an example of gene-family-specific heterogeneity, where rate changes affect the total history of specific gene families (note that in this case ultrametricity still holds) (e.g., in replicate 1, gene family 1 evolves much slower than gene family 2). Subfigure d) shows gene-by-lineage-specific heterogeneity, where changes in rate affect particular gene tree branches independently (e.g., in replicate 1, gene family 1, the ancestral A lineage evolves faster). In addition, the level of heterogeneity can also be modulated among replicates (e.g., in subfigure c) replicate 1 shows much more heterogeneity than replicate n).

#### Species-specific substitution rate heterogeneity

A rate multiplier is sampled for each branch of the species tree, modeling how different species evolve at different speeds –e.g., due to ecological conditions, metabolic rates (Martin and Palumbi 1993) or DNA repair efficiency (Britten 1986). The *alpha* parameter can be fixed or sampled de novo for each species tree (see *Distribution-driven Parameterization*).

#### Gene-family-specific substitution rate heterogeneity

A rate multiplier is sampled for each locus tree (i.e., multiplying all the branches of a given locus tree), modeling how different gene families can evolve at a different pace –e.g., due to functional constraints (Li 1997). The *alpha* parameter can be sampled independently for each species tree.

#### Gene-by-lineage-specific substitution rate heterogeneity

A rate multiplier is sampled for each branch of the gene tree, modeling how different gene tree branches evolve at different speeds due to gene-lineage interactions –e.g., due to selective bursts (Wallis 1994). The *alpha* parameter can be fixed or hierarchically sampled across the different simulation layers (e.g., the most complex parameterization would sample a hyper-hyper parameter for each species tree, a hyper parameter for each locus tree and an alpha parameter per gene tree).

The baseline substitution rate is measured in number of substitutions per site per generation. Nevertheless, *SimPhy* also takes into account absolute time units and therefore incorporates a generation time parameter. In order to increase flexibility, species-specific generation times can also be specified (see Kohne 1970).

Note that if the final objective is the simulation of multiple sequence alignments, a fifth layer of rate variation, in this case among sites (Yang 1996), can be specified using INDELible.

### Input and Output

*SimPhy* has been implemented as a non-interactive command line program. Input parameters can be given in the command line or specified in a configuration file. The program will output more or less information depending on a verbosity level controlled by the user. Output files include a species tree file, a locus tree file with all the locus trees, and one gene tree file per locus tree (typically containing a single gene tree). Depending on the settings, the program can also print out the species tree / locus tree mappings and the locus tree / gene tree mappings. Moreover, *SimPhy* can also generate an SQLite3 database with all the simulation parameters per species, locus and gene tree and some extra statistics.

## VALIDATION

We performed a series of validation experiments, described below, in order to test that *SimPhy*’s output fits different theoretical or numerical expectations. All the validation process was conducted using in-house Bash, Perl and R scripts [using APE (Paradis, Claude, et al. 2004) and phytools (Revell 2012)] available at (http://dx.doi.org/10.6084/m9.figshare.1466899). In addition, the whole simulation process was also carefully checked by hand in the debugging developing stage on several simple scenarios.

### Species tree simulation

For the SSA algorithm, we assayed the Yule process with 10 different birth rates (*λ*) (from 2.67 to 6.67 speciations/1M generations) –using 10000 replicates per level and a fixed tree origin (*t*) (1M generations)–, and compared the observed mean number of leaves with its theoretical expectation *E*(*N*(t)) = 2e^*λt*^ (Mooers, Gascuel, et al. 2012) (Fig.S1). For the BDSA, we assayed 100 combinations of 10 different birth rates (from 2.67 to 6.67 speciations/1M generations) and 10 taxa sizes (from 50 to 500 species), using 10000 replicates per combination, and compared the mean tree height with the one obtained using *TreeSim* (Stadler 2011) (Fig.S2).

### Locus tree simulation

We validated the observed average gene family size –i.e., after discarding losses and superfluous branches, 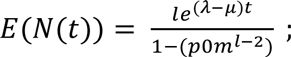; 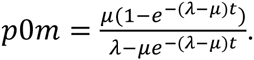. We explored 125 parameter combinations of five duplication rates (*λ*) (from 6.67 to 33.3 duplications/1E10 generations), five loss rates (*μ*) (from 0.5*λ* to 0.167*λ*) and five number of species (*l*) (from 50 to 250 species), with a fixed species tree height (*t*) (1E9 generations) and 10000 replicates per combination (Fig. S3). In addition, since gene family size is not affected by HGT or GC, we tested the expected number of horizontal gene transfer events *E*(*T*(*l*)) = *hl* under 25 parameter combinations of five different horizontal gene transfer rates (*h*) (from one to 20 transfers/1M generations) and five locus tree lengths (*l*) (from 42 to 260 M generations), with 10000 replicates per combination (Fig. S4).

### Gene tree simulation

We validated the simulation of the multispecies coalescent comparing the time to the most recent common ancestor (TMRCA) obtained with *SimPhy* and DendroPy (Sukumaran and Holder 2010) on 50 different scenarios (number of species sampled from a Uniform(50,500), speciation rate of one speciation/1M generations and effective population size sampled from a Lognormal(14,0.4)) with a total of 10000 gene trees (Fig. S5). The multilocus coalescent process was validated using a similar approach, comparing the average TMRCAs of the subtrees modeled by the bounded multispecies coalescent process obtained with *SimPhy* and DLCoal_sim (fixed species tree, five locus trees, duplication rate 0.2 duplications/1M generations and effective population size 1M individuals) with a total of 10000 gene trees (Fig. S6).

All the validation checks were clearly successful, showing only negligible deviations from the expectations or when compared to other simulations programs in spite of the large variance induced by the different evolutionary processes implemented.

## BENCHMARKING

We carried out four experiments to characterize *SimPhy*’s computational efficiency and scalability, where CPU time was defined as the sum of user and system time returned by the BSD time command. For scenarios without HGT or GC we also computed the running times of DLCoal_sim (Rasmussen and Kellis 2012). For simplicity we used a classic grid-like parameterization with 10 replicates per scenario and one individual per species. All the output options of *SimPhy* were active. Errors and running times over 300 seconds were treated as missing data. All the analyses were run in a MacBook Pro Intel Core i7 2.3Ghz, 8GB of RAM and a Solid State Drive. A generalized linear model with a Gamma error distribution was fitted to the resulting data to assess the relationship between running times and the variables studied. All the scripts used to carry out this benchmarking are available at (http://dx.doi.org/10.6084/m9.figshare.1466899).

### Benchmark 1

The first benchmark was designed to check the general performance and scalability of *SimPhy* with an increasing number of gene trees simulated under the joint effect of ILS, GDL, HGT and GC (Fig. 5). We simulated 30 different scenarios under three different schemes (#species trees/#locus trees per species tree/#gene trees per locus tree): 1/1/100-1000, 1/100-1000/1 and 100-1000/1/1. For each scenario we simulated 50-taxon species trees with a tree height of 1M generations, a speciation rate of 1E-5 speciations/generation and an effective population size of 1E4. We specified moderate but equal duplication, loss, horizontal gene transfer and gene conversion rates (5E-7 events/generation). We can immediately see in Figure 5 that *SimPhy* is extremely fast and that scales linearly with the number of trees. For example, it generates 1000 gene trees for 50 species under a complex model with ILS, GDL, HGT and GC (Fig. S7) in less than 2 seconds.

**Figure 5.**
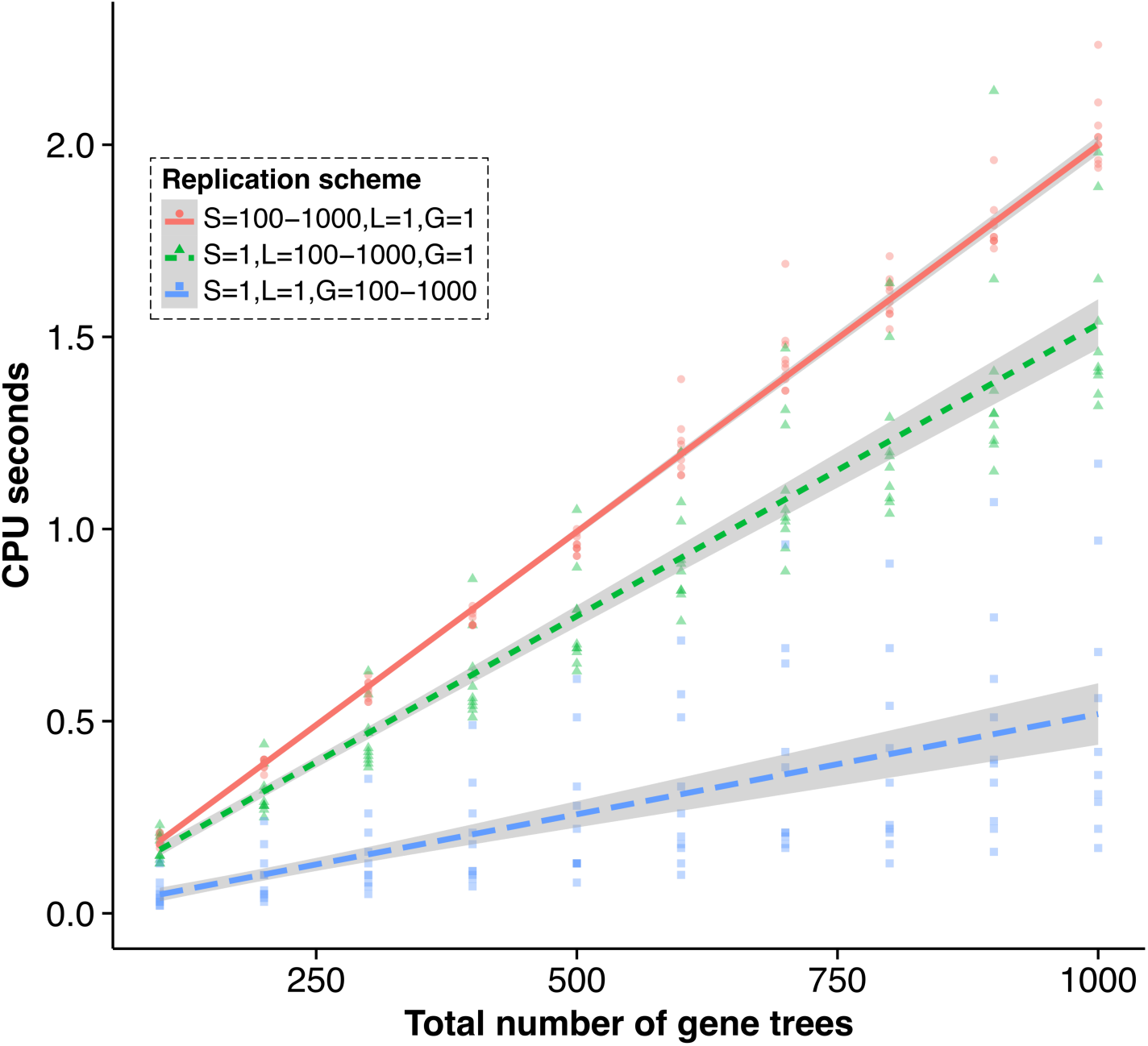
*SimPhy*’s running time against the number of gene trees simulated. A generalized linear model with a Gamma error distribution and the identity function as link was fitted to each data series. See *Benchmark 1* for the simulation details.

### Benchmark 2

This benchmark was intended to compare *SimPhy* and DLCoal_sim under a very simple model, with only moderate levels of ILS. We designed 10 scenarios with 10 numbers of species, varying from 50 to 500. For each scenario we simulated a single species tree, assuming a speciation rate of 1E-6 speciations/generation an effective population size of 1E4, and generated 100 gene families. In Figure S8 we can see that *SimPhy* scales linearly with the number species, while DLCoal_sim only does the same for small species trees, being unable to complete simulations with more than 300 species. Remarkably, *SimPhy* was at least one order of magnitude faster than DLCoal_sim. This might be explained by the fact that *SimPhy* differentiates subtrees that can be modeled by the multispecies coalescent from those that require the bounded alternative, using a much faster and less error prone algorithm for the former situation. Moreover, in the absence of GDL, HGT or GC locus trees and species trees are equivalent, so in these cases *SimPhy* directly uses the species trees as locus trees instead of simulating them *de novo*. In addition, *SimPhy* and *DLCoal_sim* also have important implementation differences –e.g., data structures, coding language– which make *SimPhy* faster in most scenarios. Finally, *SimPhy* uses a multiple precision library for very complex scenarios to favor its scalability.

### Benchmark 3

In this case we characterized *SimPhy*’s and DLCoal_sim’s running times for a model with moderate levels of ILS and GDL. We designed 30 scenarios, with 100, 200 and 300 species and 10 duplication rates –from 1E-6 to 3.25E-6 duplications/generation. For each scenario we simulated a single species tree, assuming a speciation rate of 1E-5 speciations/generation an effective population size of 1E4, and generated 100 gene families. Logically, duplications increased the running time for both programs, but with *SimPhy* still being much faster than *DLCoal_sim* (Fig. S9). Moreover, *DLCoal_sim* showed again important scalability problems, being unable to simulate trees with 300 species.

### Benchmark 4

Here we aimed to evaluate the sampling efficiency of the multilocus coalescent model by exploring a model with various levels of GLD and ILS. We designed 30 scenarios with 10 effective population sizes –from 976 to 5E5 individuals– and three duplication rates –0, 1E-6 and 2E-6 duplications/generation. For each scenario we simulated a single 100-taxon species tree, assuming a speciation rate of 1E-5 speciations/generation and a tree height of 5E5 generations, and generated 100 gene families. For both programs we observed an important increase in execution time with higher ILS in the presence of duplications, and almost no effect in their absence (Fig. 6). Importantly, *SimPhy* was again at least one order of magnitude faster and scales much better than *DLCoal_sim*. Several *DLCoal_sim* replicates were allowed to run beyond the 300-second limit but did not finished even after 24 hours. This behavior is likely reflecting a problem in the rejection-sampling algorithm that *DLCoal_sim* uses to sample the bounded multispecies coalescent (specifically the lineage counts), since this algorithm is prone to get stuck sampling highly improbable scenarios. On the other hand, *SimPhy* uses an alternative sampling strategy (see *Simulation Process* and Appendix 1) that seems to effectively mitigate this problem.

**Figure 6.**
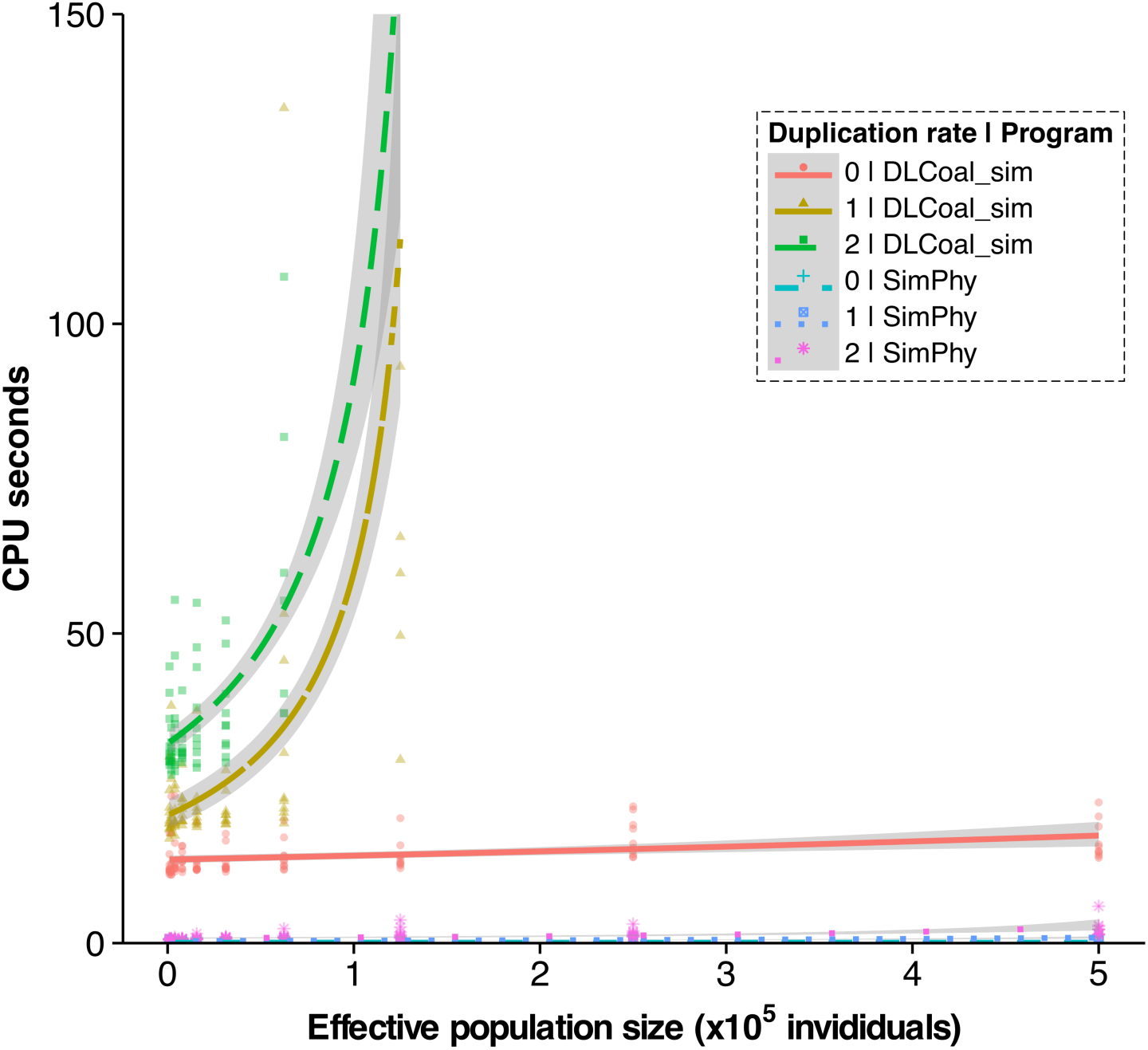
Running time comparison between *SimPhy* and DLCoal_sim under a GDL+ILS model. Scatter plot of the time needed by the two simulators to generate 100 gene trees from 100 locus trees using three different duplication rates (duplications/1M generations) under different effective population sizes. A generalized linear model with a Gamma error distribution and the inverse function as link was fitted to each data series. DLCoal_sim’s execution times over 150 seconds are not shown.

## TEST-CASE STUDY: DUPLICATION TIME OVERESTIMATION OF DL RECONCILIATIONS

We have previously shown that the most recent common ancestor (MRCA) of a new gene originated by duplication and its paralog does not necessarily coincide with the individual where this duplication first occurred, generating a systematic overestimation of the duplication time for locus-tree unaware reconciliation methods (Mallo, De Oliveira Martins, et al. 2014b) (Fig. S10). Here we further explored this issue by quantifying this overestimation in more complex scenarios for which theoretical expectations are not available. In order to do so, we simulated 1E6 replicates with a number of species uniformly distributed from 25 to 200, with a fixed speciation rate (1E-5 speciations per time unit) an effective population size uniformly distributed from 1E3 to 10E3 individuals, and with just one gene family per genome. The locus tree simulation was parameterized with a duplication rate sampled from a Uniform distribution of 5E-8 to 5E-7 events per generation, and with a number of individuals per species uniformly ranging from one to five. To check the relationship between the simulated parameters and the overestimation bias, we performed a stepwise selection under the AIC criteria of a generalized linear model with Gamma error distribution and inverse link on a linear combination of all the simulation parameters. As dependent variable we used the mean of the distance between the real duplication node (locus tree) and the first coalescence between lineages coming from the two different paralogs for all the duplications for each simulation replicate.

Our results show that the effective population size increases the overestimation bias, while the number of individuals per species and the duplication rate have the opposite effect, being the only three parameters retained in the best-fit linear model. That a larger effective population size increases the overestimation bias is expected, as the coalescent time of any two lineages grows with it. An increasing number of individuals per species should reduce this bias, as with more lineages going through the duplication node the time for the first coalescence among paralogs decreases. The effect of the duplication rate, inversely correlated with this bias, is less clear. Nevertheless, it can be tentatively explained by the coalescent bounds imposed by duplications. Expected coalescent times along a bounded subtree are shorter than along an unbounded subtree, since the probability of coalescence is scaled by the probability of the TMRCA being less or equal the bound (i.e., we can think of the unbounded coalescent as a bounded coalescent with the bound in the infinite, and therefore scaled by one). Thus, the bigger the duplication rate, the bigger the probability of having duplications in bounded subtrees, and the bigger the reduction on the expected duplication time overestimation generated by the coalescent bounds.

Interestingly, we can conclude that the expected overestimation for the simplest case – 2Ne for one diploid individual with two paralogs; (Mallo, De Oliveira Martins, et al. 2014b)– constitutes an upper bound, with a smaller bias expected in more complex scenarios (with duplications and multiple individuals per species). Finally, we also note that the size of the species tree did not seem to play a significant role in this case.

## CONCLUSION

We have introduced, validated and benchmarked *SimPhy*, to our knowledge the first software that simulates gene tree family evolution under the three main evolutionary processes that generates species tree/gene tree incongruence –ILS, GDL and HGT. This, together with its comprehensive heterogeneity models and the parameter sampling strategy should help *SimPhy* becoming a powerful phylogenomic tool. We have also conducted a simple case study in order to show a potential application (duplication time overestimation), but we envision many more to come. In fact, *SimPhy* has been already used to validate at least three species tree reconstruction methods (De Oliveira Martins, Mallo, et al. 2014; Bayzid, Mirarab, et al. 2015; Mirarab and Warnow 2015).

## AVAILABILITY

*SimPhy* is distributed under the license GNU GPL v3. It is written in C and it relies on four libraries, the GNU Scientific Library (GSL), the GNU Multiple Precision Arithmetic Library (GMP), the GNU MPFR Library and SQLite3. Users can find the source code, pre-compiled executables, a detailed manual and example cases on a GitHub repository (https://github.com/adamallo/SimPhy). We provide two flavors of pre-compiled executables, for Linux –Linux Standard Base compatibility (73 out of 74 for the 64 bits version, 84 out of 87 for the 32 bits version)– and MacOSX systems.

## ACKNOWLEDGMENTS

We want to thank Tanja Stadler for her kind help in the early-stage development of *SimPhy* (birth-death process sampling algorithms).

## Appendix 1 Lineage counts sampling strategy SimPhy: Phylogenomic Simulation of Gene, Locus and Species Trees

In order to sample the bounded multispecies coalescent *SimPhy* uses a three-step strategy similar to the one used by DLCoal_sim (Rasmussen and Kellis, 2012):

i. Calculation of the probabilities of the number of lineages (counts) going into and out of every locus tree branch of the considered tree/subtree, performed backwards in time.
ii. Sampling of the lineage counts using a forward in time traversal.
iii. Sampling of coalescent times conditioned on the just obtained counts in an additional – backwards in time– traversal.

While for steps i) and iii) *SimPhy* uses its own implementation of the original DLcoal_sim algorithms, the original algorithm for sampling linage counts has been modified in order to improve the scalability of the simulations, changing the original rejection sampling algorithm for an inverse transform sampling algorithm. This alternative sampling method is applied on the the cumulative distribution function of the number of input lineages *a*(*u*) entering the branch *e*(*u*) of the considered locus tree node *u* conditioned on the output lineage counts *b*(*u*) and locus tree parameters *θ* = (*T*^*L*^, **t**^*L*^, **N**, **n**) (topology *T*^*L*^, branch lengths **t**^*L*^, population sizes **N** and gene tree copies per locus leaf **n**). The number of input lineages *a*(*u*) is afterwards distributed –taking into account the count probabilities– between the descendants *c*_1_(*u*) and *c*_2_(*u*) to become its new output node counts *b*(*c*_1_(*u*)) and *b*(*c*_2_(*u*)). In order to devise this function, let us start by the probability distribution of the number of lineage counts entering a locus tree lineage, given the output lineage counts and trees (Rasmussen and Kellis, 2012):

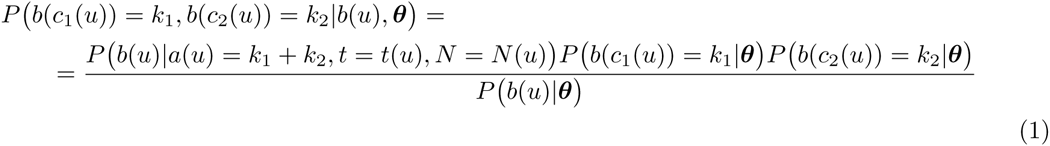

Focusing on sampling the input counts *a*(*u*), let us consider a random variable *k*_*t*_ = *b*(*c*_1_(*u*)) +*b*(*c*_2_(*u*)). Consequently, we have the probability function:

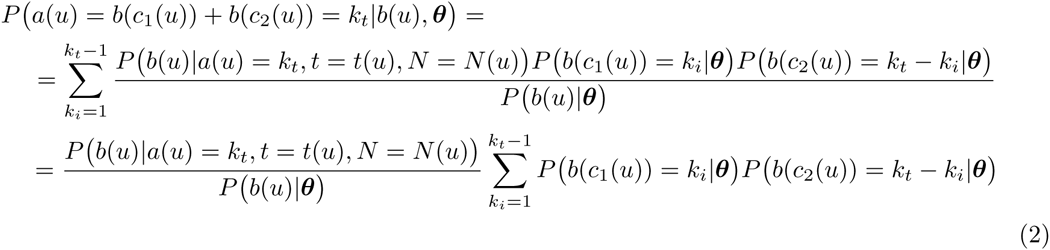

Now, having *P*(*a*(*u*) = *k*_*t*_ |*b*(*u*), ***θ***) we can obtain the target CDF, taking into account that *k*_*t*_ is a discrete random variable:

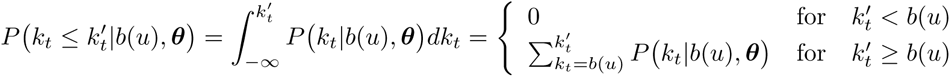

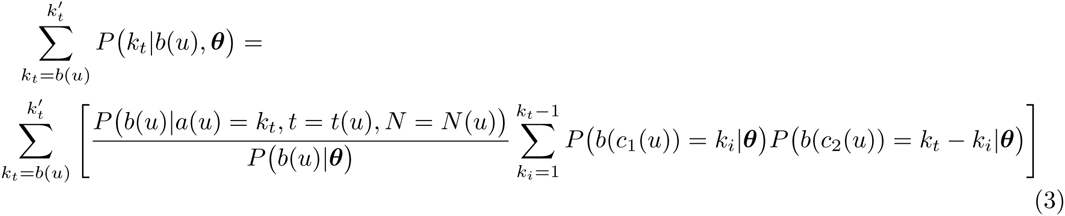

## Supplementary Material

**Fig. S1.**
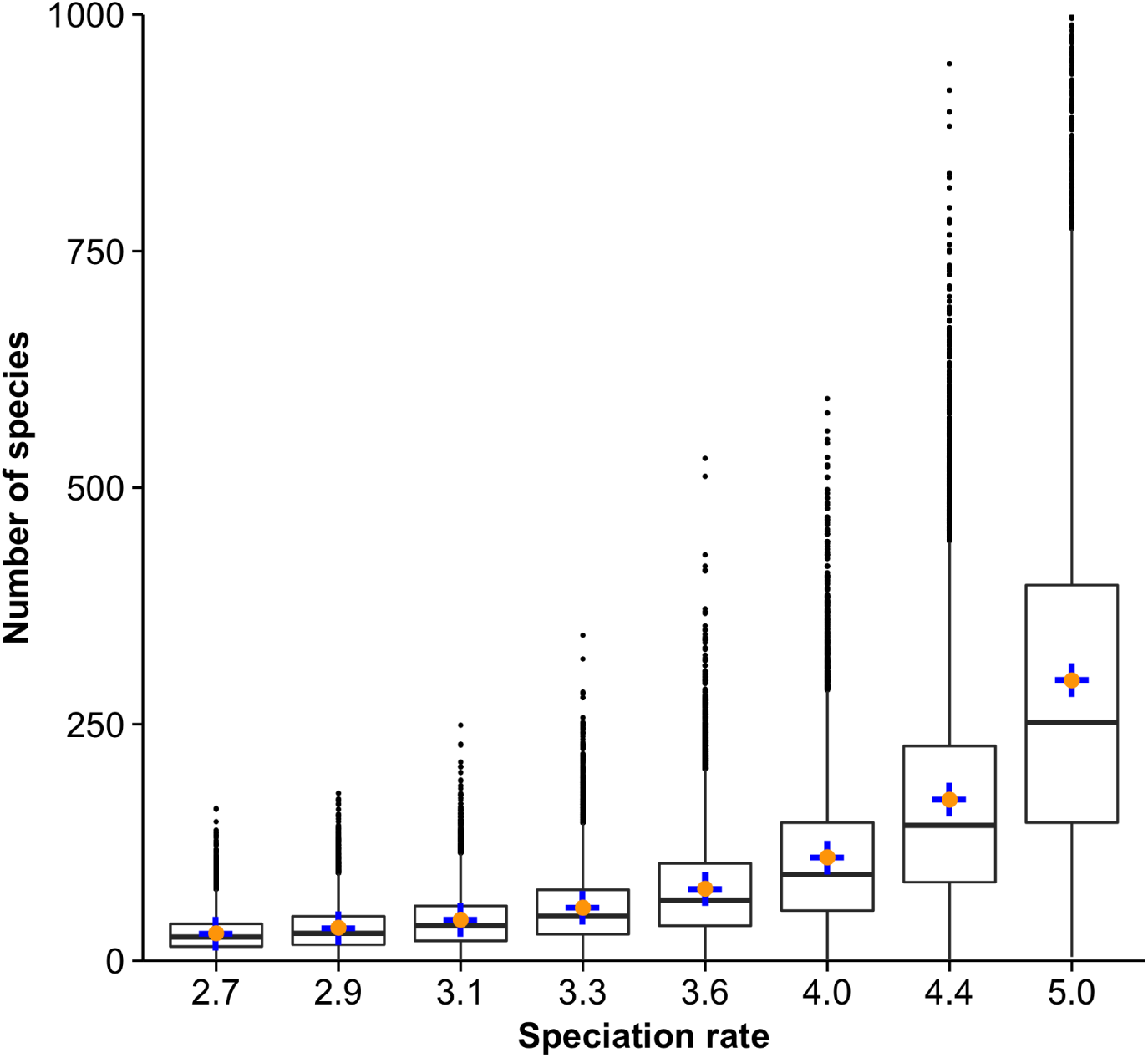
Validation of *SimPhy*’s species tree simulation (variable number of species). Boxplots describe the distribution of the number of species generated by 10000 simulation replicates across different speciation rates (speciations/1M generations) given a fixed tree height (1M generations). Expected theoretical values are indicated with a blue cross, while the observed average values are depicted with an orange dot. For representation purposes, extreme values in the rightmost boxplot are not shown.

**Fig. S2.**
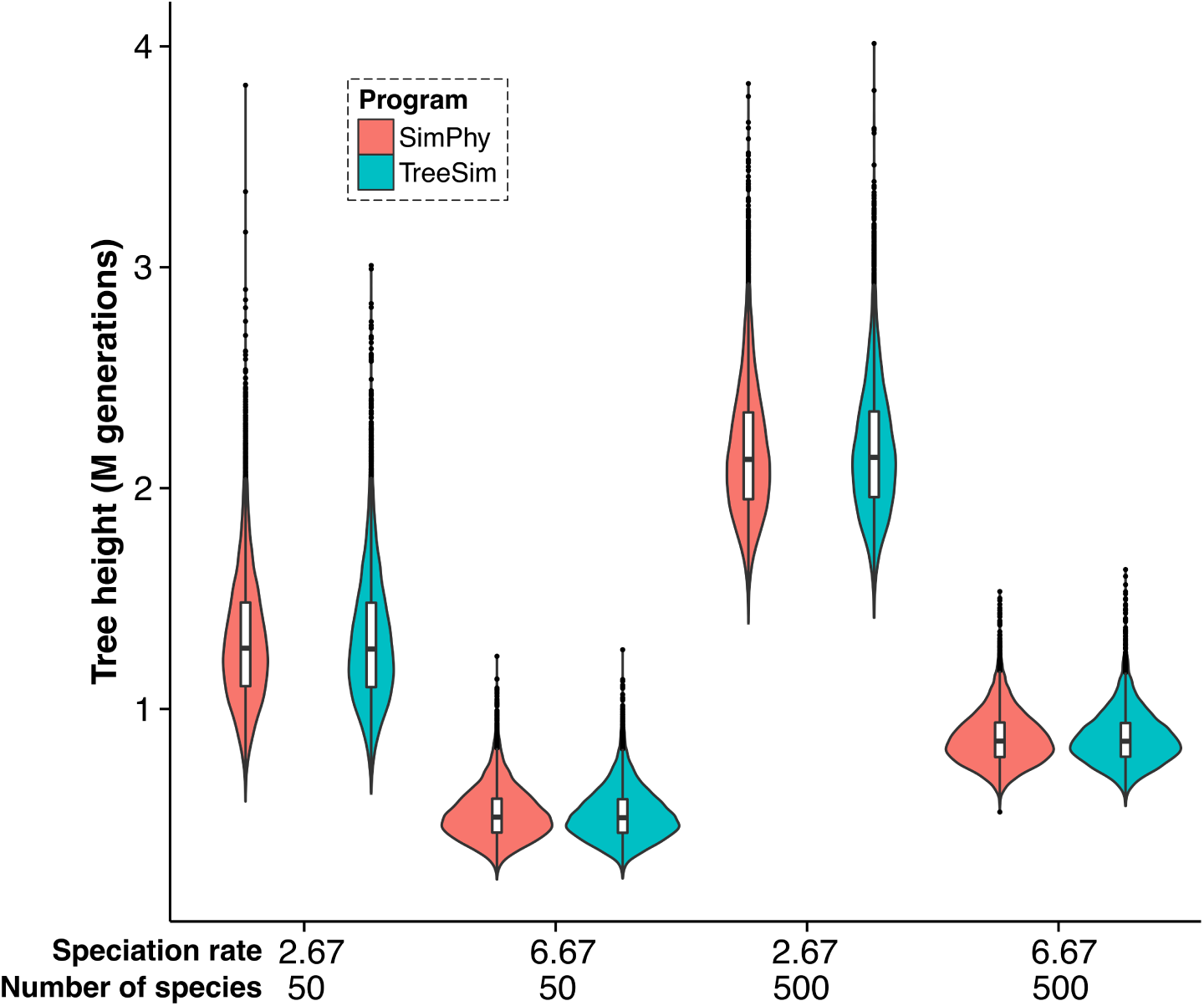
Validation of *SimPhy*’s species tree simulation (variable tree height). Violin plots (kernel density curve with a boxplot inside) describe the distribution of 10000 species tree heights simulated by *SimPhy* (red) and TreeSim (blue) (speciations/1M generations).

**Fig. S3.**
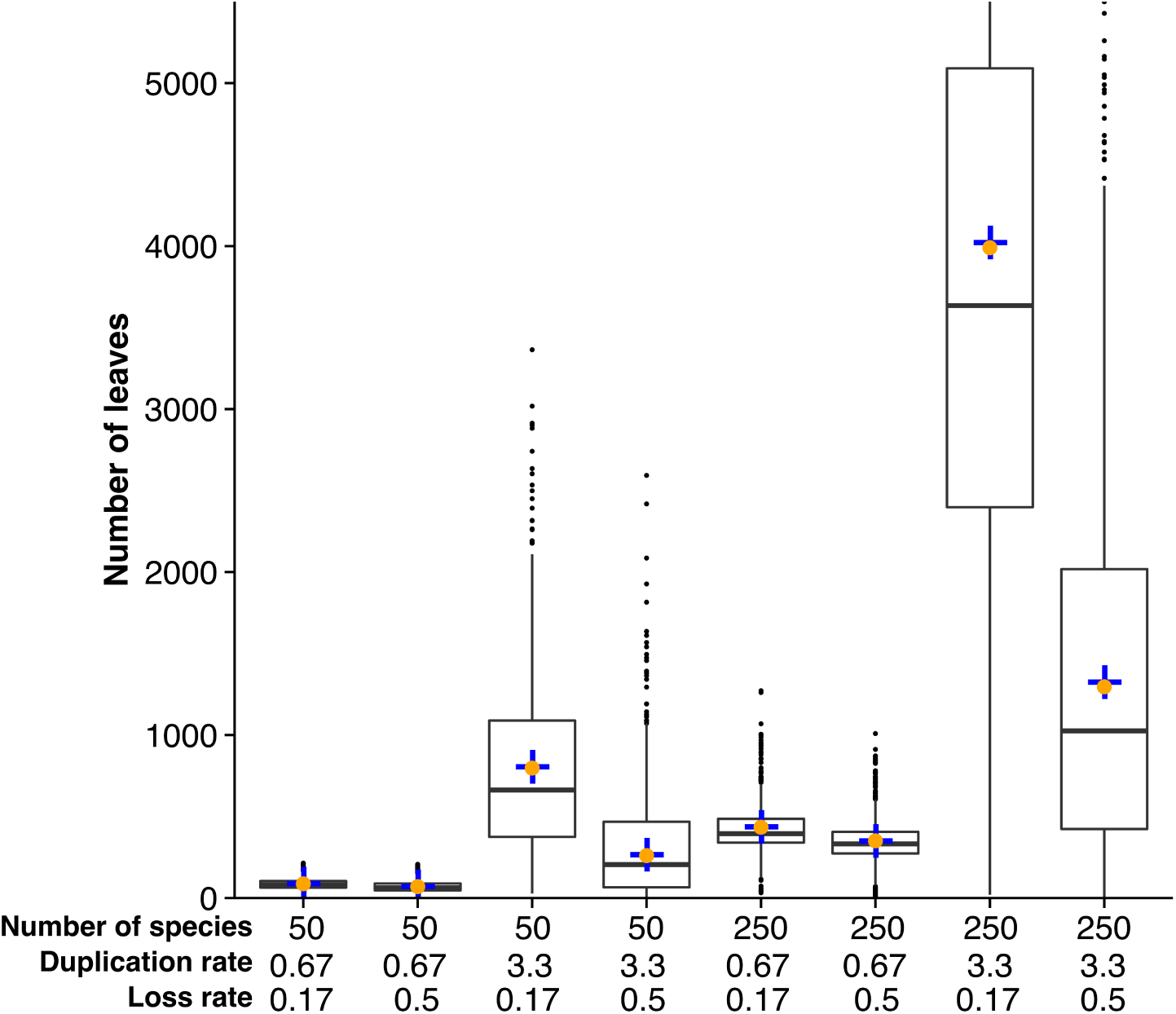
Validation of *SimPhy*’s locus tree simulation under a GDL model. Boxplots describe the distribution of the number of locus tree leaves generated by 10000 simulation replicates across different duplication rates (speciations/1M generations), loss rates (relative to the duplication rate) and number of species. Expected theoretical values are indicated with a blue cross, while the observed average values are depicted with an orange dot. For representation purposes, some extreme values are not shown.

**Fig. S4.**
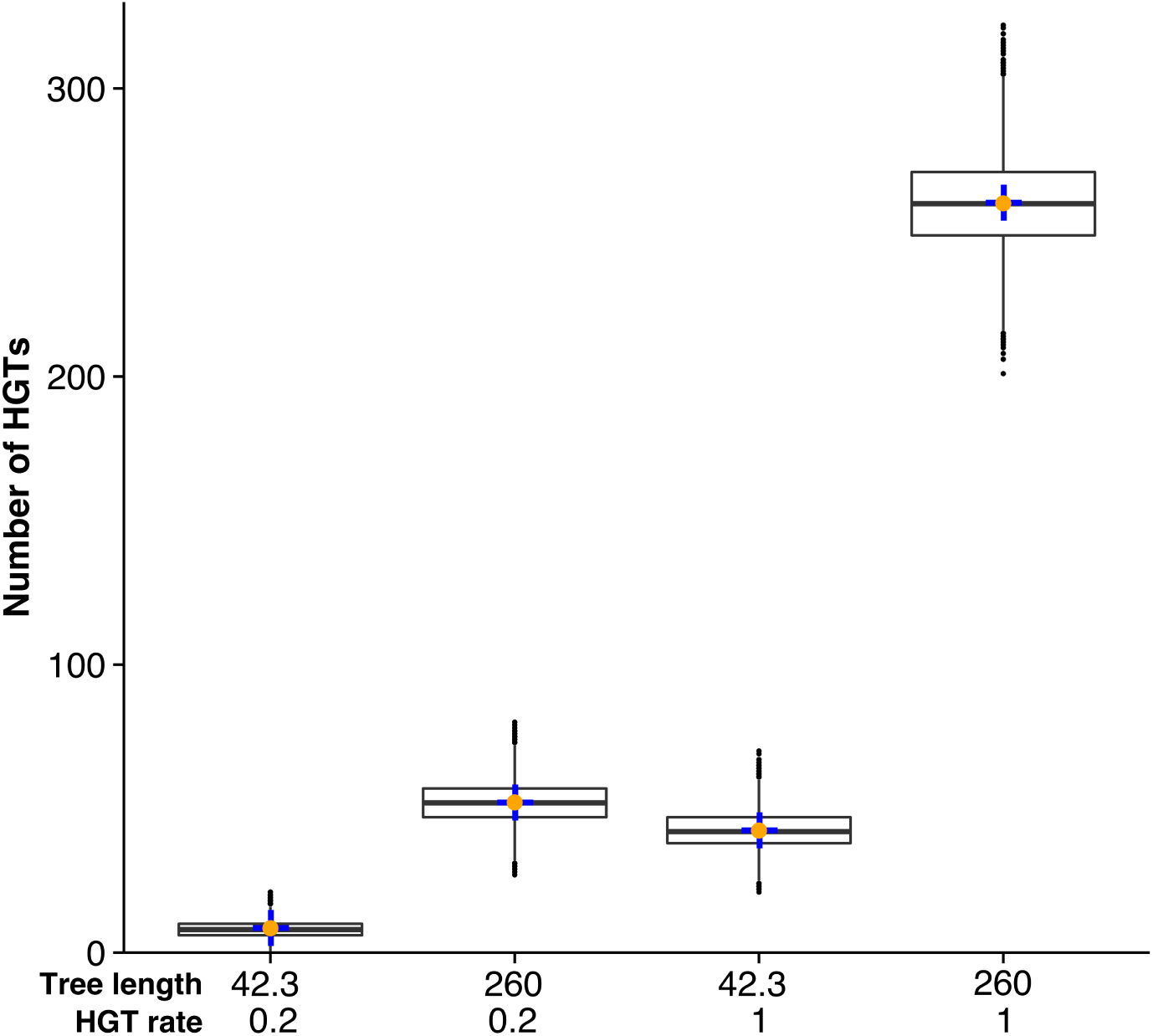
Validation of *SimPhy*’s locus tree simulation under an HGT model. Boxplots describe the distribution of the number of HGT events per locus tree generated by 10000 simulation replicates across different HGT rates (transfer/1M generations), and tree lengths (M generations). Expected theoretical values are indicated with a blue cross, while the observed average values are depicted with an orange dot.

**Fig. S5.**
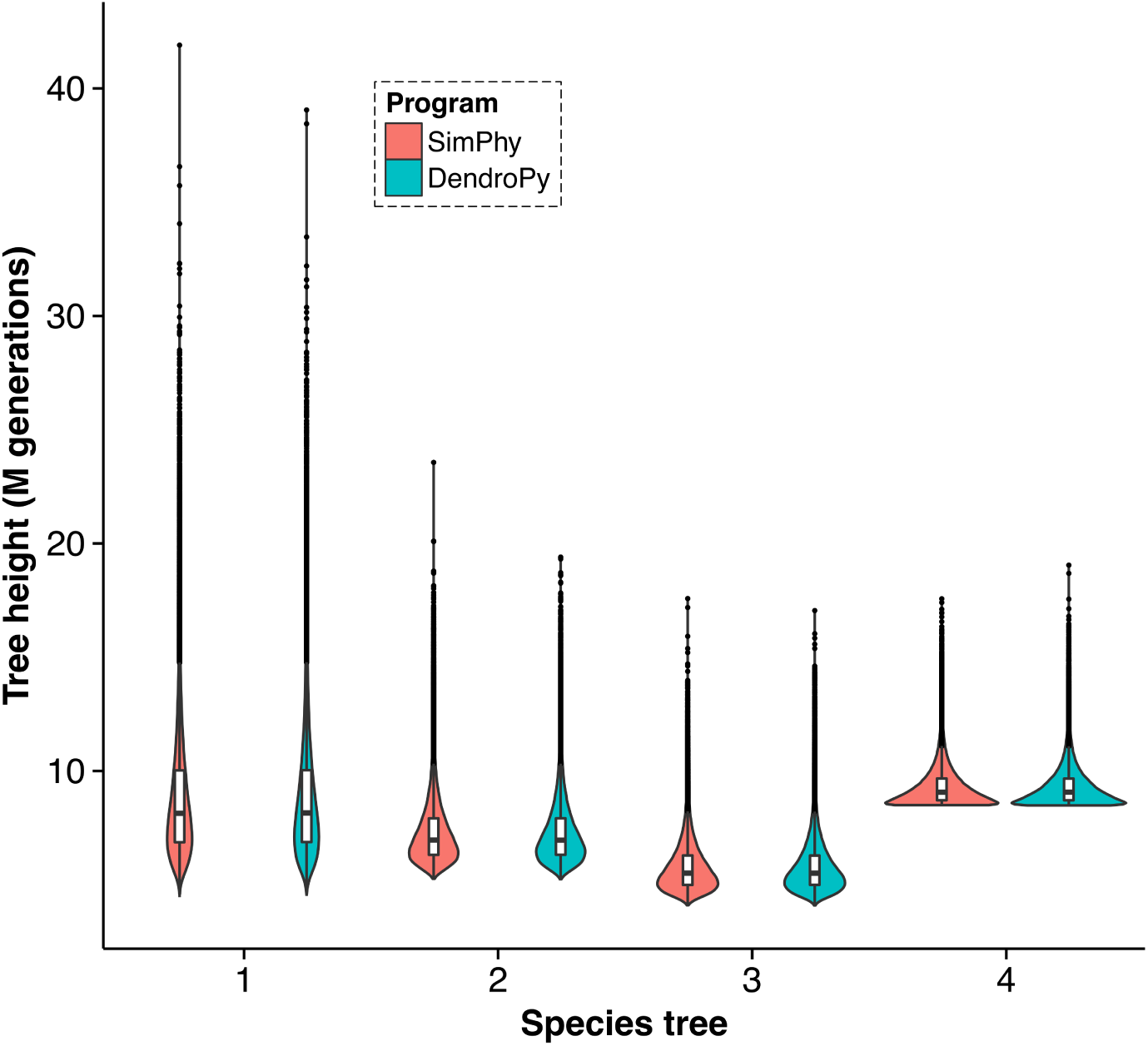
Validation of *SimPhy*’s bounded multispecies coalescent sampling. Violin plots describe the distribution of 10000 gene tree heights simulated by *SimPhy* (red) and DendroPy (blue) under four different species trees.

**Fig. S6.**
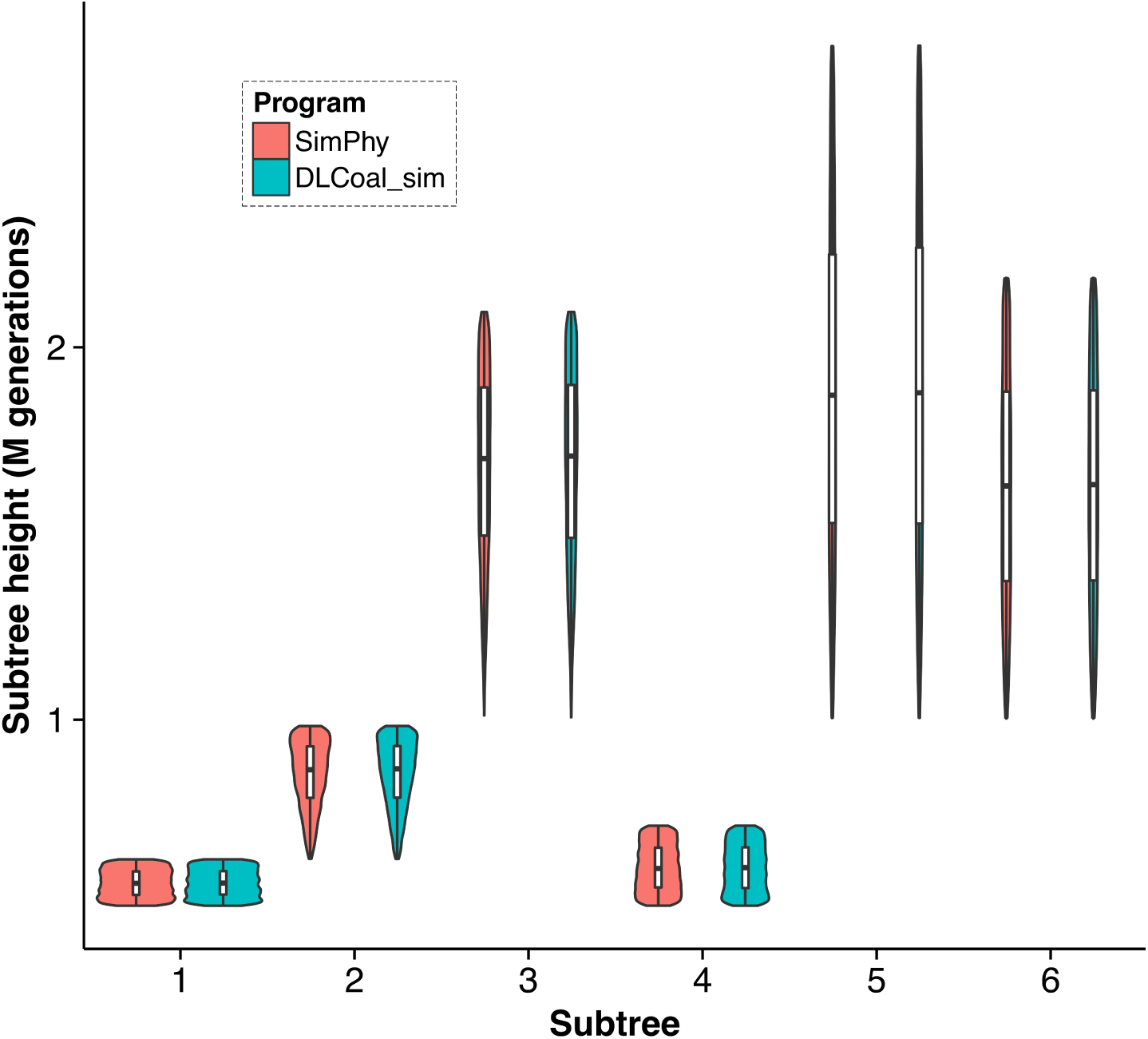
Validation of *SimPhy*’s bounded multilocus coalescent sampling. Violin plots describe the distribution of 10000 gene subtree heights simulated by *SimPhy* (red) and DLCoal_sim (blue) under six different bounded locus subtrees.

**Fig. S7.**
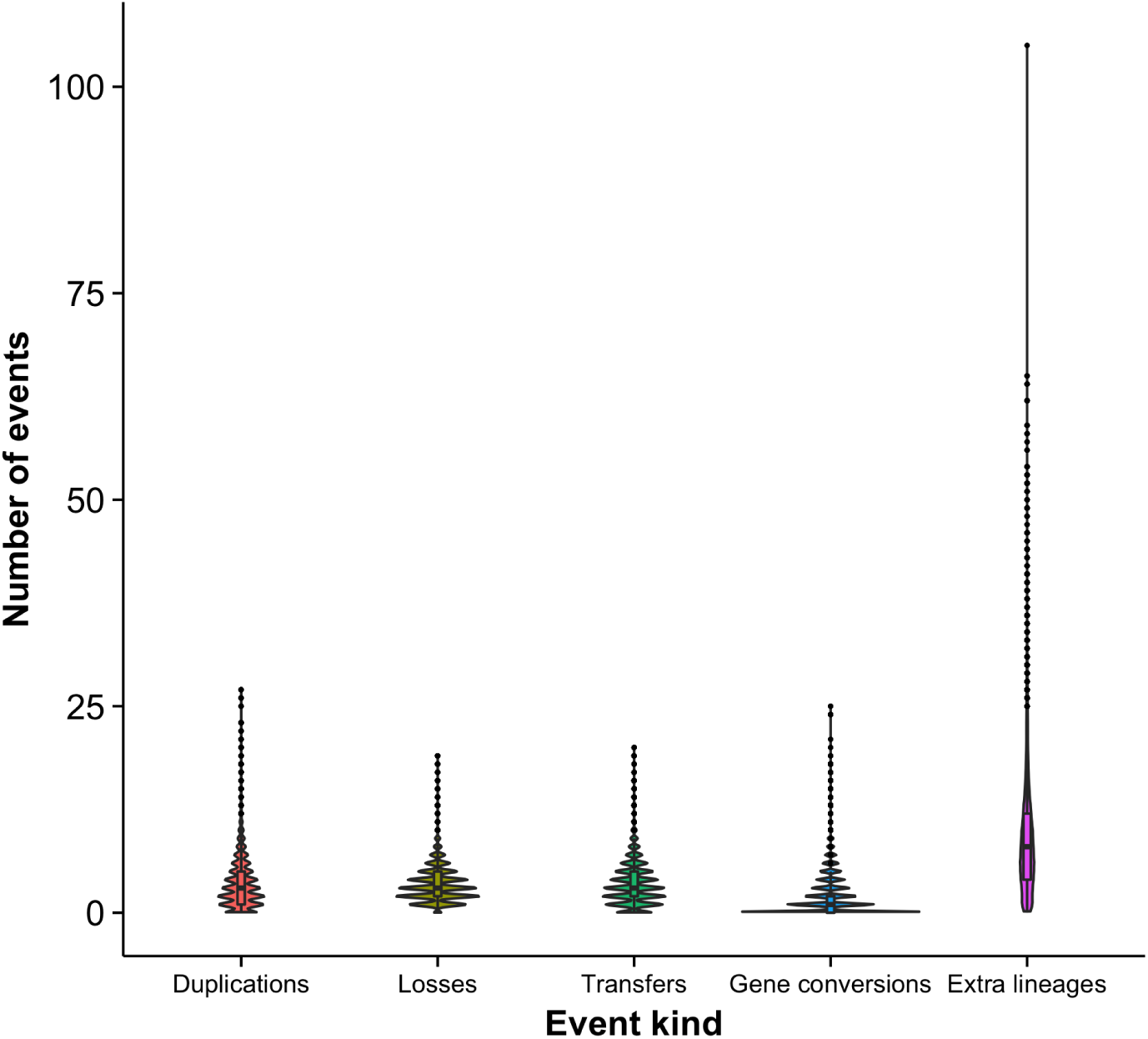
Distribution of the number of evolutionary events in benchmark 1. Each violin plot represents the distribution of the number of events per tree.

**Fig. S8.**
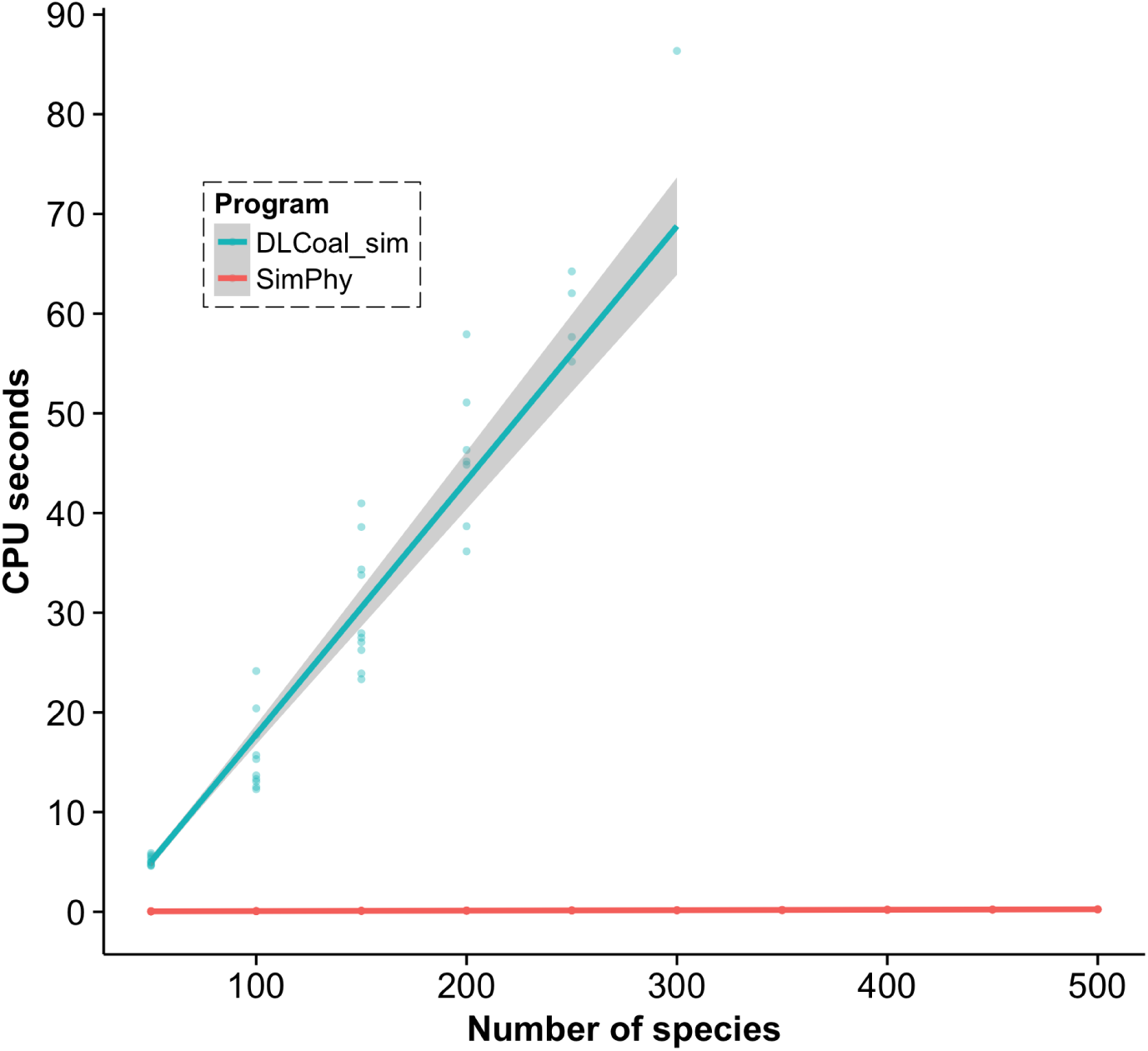
Running time comparison between *SimPhy* (red) and DLCoal_sim (blue) under an ILS model. One hundred gene trees were simulated along 100 locus trees with different numbers of species. A generalized linear model with a Gamma error distribution and the identity function as link was fitted to each data series. DLCoal_sim was unable to run with more than 300 species. Note that the execution time of *SimPhy* also includes the species tree simulation, while DLCoal_sims execution time does not.

**Fig. S9.**
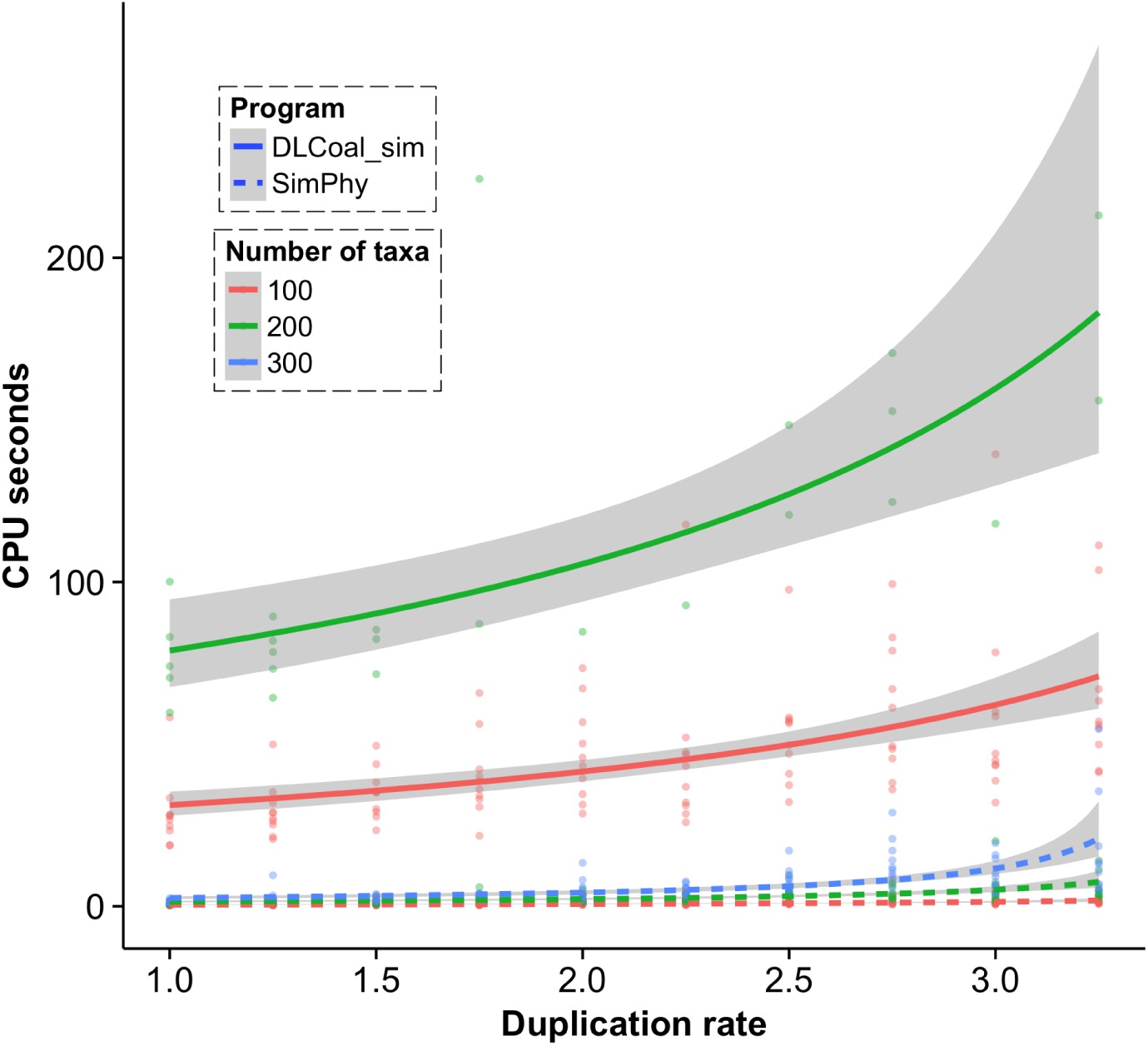
Running time comparison between *SimPhy* (dashed lines) and DL-Coal_sim (solid lines) under an GDL+ILS model. One hundred gene trees were simulated along 100 locus trees with different numbers of species and duplication rates (duplications/1M generations). A generalized linear model with a Gamma error distribution and the identity function as link was fitted to each data series. DLCoal_sim was unable to run with more than 300 species.

**Fig. S10.**
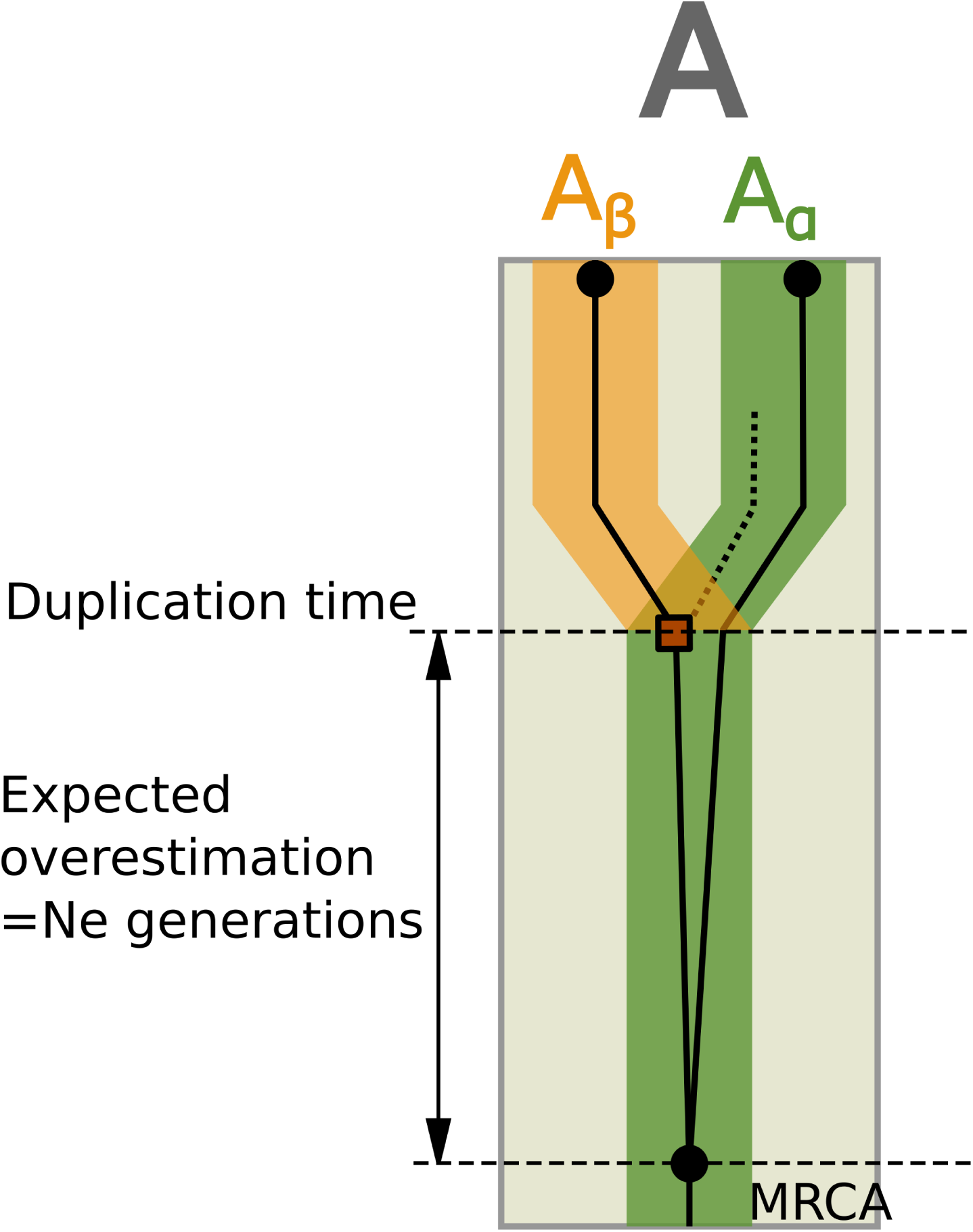
Overestimation of the duplication time. Here we depict the overestimation on the duplication time for two paralogous copies (*A*_*α*_, *A*_*β*_) in a single individual (*A*). The gene tree (thin black lines) evolves inside the locus tree (medium-thick green and orange lines) along a species tree branch (light gray shadow in the background). The duplication takes place in the gene copy represented by a red square, whose lineage in the green locus does not reach the present (dashed line). For haploid populations, the expected overestimation is Ne generations.

